# Towards One-Step and Universal Synthesis of Hypermodified Uracil Nucleosides: acp^3^U and cmnm^5^U

**DOI:** 10.1101/2025.09.26.678795

**Authors:** Ewa Mejdr, Lena Heinickel, Thomas Carell, Ivana Mejdr

## Abstract

Hypermodified nucleosides have been recently in the center of scientific attention. They represent a unique group of nucleosides with an alternated structure, such as the addition of functional groups or amino acids. Their distinctive structures and positions in RNAs are crucial for the processes of translation and stability. High demand of oligonucleotides bearing those hypermodified nucleobases lead us to the development of a single step synthesis of acp^3^U phosphoramidite, as acp^3^U has been recognized as an important molecule for the structural integrity of tRNA and native immunity. We also present a novel synthesis of cmnm^5^U phosphoramidite and its incorporation into an oligonucleotide from a universal starting material, allowing a transformation into at least two other hypermodified nucleosides.

## Introduction

The study of hypermodified nucleosides has been a hot topic for some years now. Various types of RNAs are enriched with non-canonical nucleosides that differ structurally from the canonical nucleotides to a significant extent.^1, 2^ Hypermodified nucleosides cover a broad range of functions in RNA molecules, such as structural stabilization to aiding the decoding process. Up to date around 150 modifications have been identified in RNAs from all realms of life.^1, 2^ The transfer RNA (tRNA) relies especially on those modifications that help to stabilize the tertiary structure or tune the decoding proficiency and guarantee a proper translation. A wide range of modifications are located in the region of the anticodon loop, where they modulate the codon-anticodon interaction whereas others could be found in the D-loop and T-loop, where they have a positive influence on correct folding. Hypermodification of all four canonical nucleosides has been observed. In our previous works, we have focused on amino-acid decorated nucleosides like t^6^A or m^6^t^6^A and C5-modified uridine structures like nm^5^U or mnm^5^U, found at positions 37 and 34 of the anticodon loop of tRNA, respectively. It has been suggested that some of the modified nucleosides can be considered ‘living molecular fossils’,^3^ relics of an ancient RNA-world.^2, 4, 5^

These anticodon modifications have captured our attention in the context of the origin of life research. We have postulated a concept of an RNA-peptide world, where amino acid decorated nucleosides^6^ in concert with (m)nm^5^U modification play a crucial role.^7^ We have shown the possibility of amino acid transfer between the modifications and the stereoselectivity thereof.^8,9^

We are currently investigating in particular on two uracil modifications 3-(3-amino-3-carboxypropyl)uridine (acp^3^U) and 5-carboxymethylaminomethyluridine (cmnm^5^U), which are derived from the amino acids homoserine and glycine, respectively.

acp^3^U, first discovered in the tRNA of Escherichia coli^10^, is a highly conserved modification found in the variable loop and D-loop of tRNAs at positions 47 of bacterial and at position 20 and 20a in eukaryotic tRNAs, as well as in archaeal rRNA.^11^ It has been identified in a signature sequence of prokaryotic and eukaryotic tRNAs^12^ and plays a crucial role in maintaining tRNA structural integrity by preventing unwanted base pair interactions.^13, 14^ The acp group is attached to the N3 position of the uracil base, effectively inhibiting its participation in Watson–Crick base pairing.^13^ Despite its widespread conservation across bacterial and eukaryotic tRNAs, the precise mechanisms underlying the biogenesis and physiological functions of acp^3^U remain unclear. However, studies recognized TapT enzyme (tRNA aminocarboxypropyltransferase) in E. coli as well as human homologs DTWD1 and DTWD2 as responsible for the acp^3^U formation in tRNA.

Recently, it has been reported that in glycoRNA, the acp^3^U modified nucleoside could be an attachment site for N-glycans^15^ and that N-glycans attached via acp^3^U prevent innate immunity sensing of endogenous small RNAs. Additionally, deglycosylation of acp3U modified small RNA triggered inflammatory responses in Toll-like receptor signalling pathways.^15, 16^ This finding significantly expands the functional repertoire of acp^3^U beyond its traditional role as a tRNA modification, suggesting its involvement in post-transcriptional modifications and cellular signaling pathways.^15, 17^ Glycosylated RNAs may represent an unprecedented interface between RNA biology and glycan-mediated cell communication, opening new ways for exploring RNA functionality and regulatory mechanisms.^18^

Next to acp^3^U, cmnm^5^U is a uridine modification that plays a critical role in RNA functionality. cmnm^5^U is a hypermodified nucleoside present at the wobble position in the anticodon stem loop of bacterial and eukaryotic mitochondrial tRNA.^11^ This modification is crucial for accurate codon recognition during translation by restricting the wobble base pairing, thereby enhancing the fidelity of protein synthesis.^19^ In mitochondria, cmnm^5^U is particularly responsible for the correct translation of codons ending a with purine that would otherwise be ambiguous, such as the UGA codon, which is reassigned from a stop codon to code for tryptophan in many mitochondrial genomes.^20^ The presence of cmnm^5^U at the wobble position plays an indispensable role in structural stabilization of the anticodon loop through 5′-ribose-phosphate backbone and base mediated interactions, in order to enhance the ribosome-tRNA binding ability and is preventing misreading and ensuring proper translation^19, 20^ A related modification, nm^5^U, can arise from cmnm^5^U through enzymatic decarboxylation. This base is also found at the wobble position in various tRNAs.^21^ nm^5^U lacks the carboxymethyl group present in cmnm^5^U, which may result in reduced hydrogen bonding potential and slightly altered codon recognition properties.^22^ Still nm^5^U is modulating the flexibility of the anticodon loop and supporting efficient decoding of NNA and NNG codons.^23^ Its presence is often associated with organism- or tissue-specific variations in mitochondrial translation and may reflect adaptive regulatory mechanisms.^23^ The lack of xm^5^U (x = mn, n, cmn) in human mitochondria is observed in patients with mitochondrial encephalomyopathy, lactic acidosis, stroke-like episodes and myoclonus epilepsy with ragged-red fibers (MERRF).^24^

In biological systems, the cmnm^5^U moiety is formed post-transcriptionally from uridine via an enzymatic reaction with glycine.^24^

Interestingly, cmnm^5^U is able to base pair with A or G in the codon of the mRNA,^20^ while acp^3^U is unable to undergo Watson-Crick base pairing due to the attachment of the 3-amino-3-carboxypropyl group to the N3 of uracil base.^13^

## Results and Discussion

We has previously published a six-step synthesis route for the acp^3^U phosphoramidite from uridine.^25^ Here we propose a one-step synthesis, from the uridine phosphoramidite via a Mitsunobu type reaction with excellent yield in both milligram and gram scale synthesis.

**Scheme 1:**
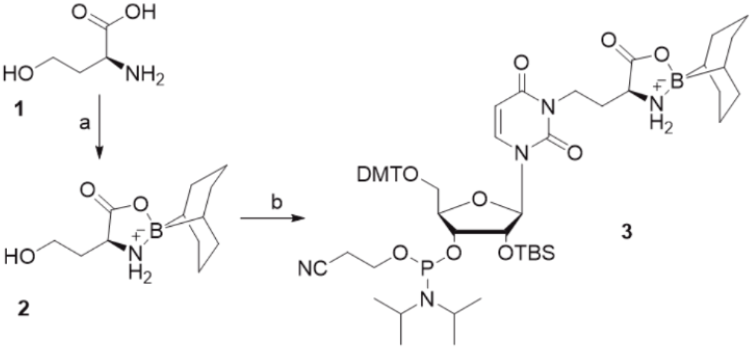
*Synthesis of acp*^*3*^*U phosphoramidite* ***3***. Reagents and conditions: a) 9-BBN, MeOH, THF, 80 %; b) 2′-OTBS uracil phosphoramidite, PPh^3^, DIAD, 1,4-dioxane, 0 °C-r.t., 3h, 92 %.

The first step of our new synthesis involves preparation of the protected L-homoserine (with 9-BBN) **2** as described previously by Nainyte et al (Scheme 1).^25^ Commercially available 5’-DMT and 2’-TBS protected uridine phosphoramidite was then used as a starting material for a Mitsunobu type reaction with compound **2**, to yield the acp^3^U phosphoramidite **3** directly. We screened the conditions in milligram scale reactions, varying the solvent and amount of BBN-protected homoserine and PPh_3_. The highest conversion and isolated yield were observed in 1,4-dioxane with 1.2 eq of reagents, where we isolated the product in 92 % yield. We then scaled up to a 2-gram scale reaction and isolated the product **3** with full separation of both formed isomers. This we could also confirm by two separate signals in ^31^P NMR measurement (Fig. 2). Similar results were obtained, although with slightly lower yields, in THF (68 %).

**Figure 1.**
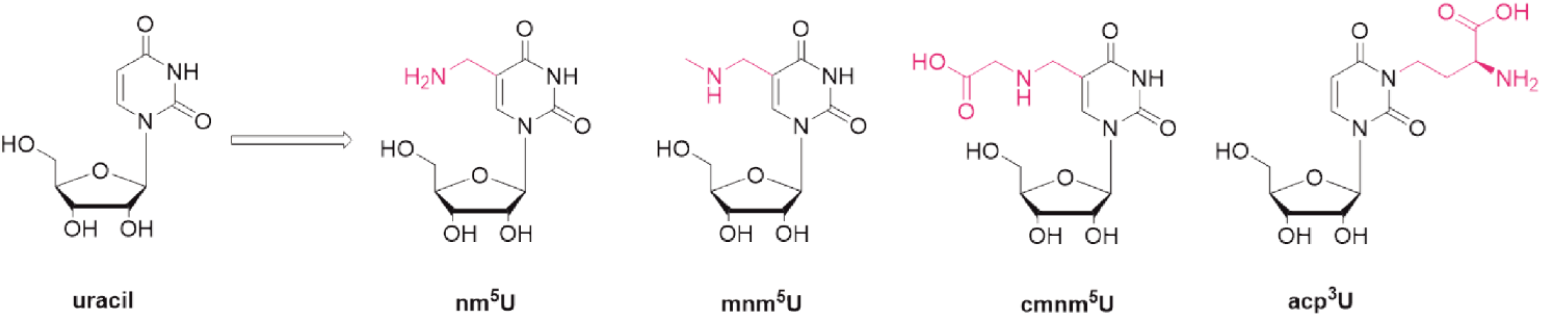
Naturally occurring hypermodified uracil nucleosides nm^5^U, mnm^5^U, cmnm^5^U and acp^3^U.

**Figure 2.**
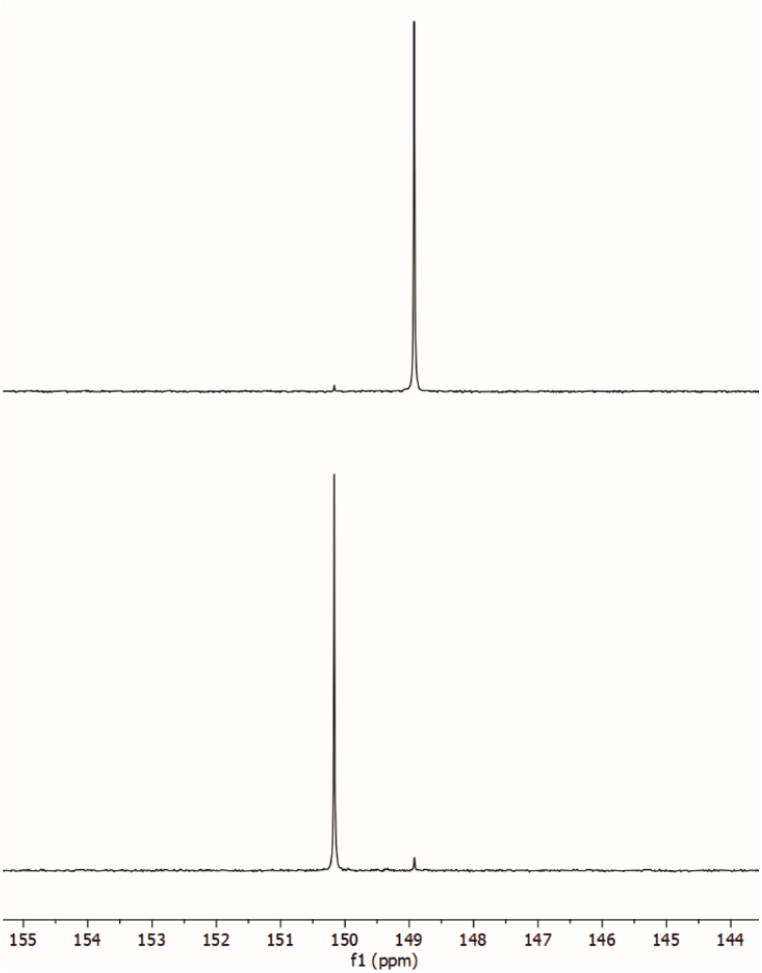
Comparison of ^31^P NMR spectra of two separated acp^3^U phosphoramidite isomers isolated from scaled-up 2 g reaction.

Next, we focused on the synthesis of cmnm^5^U. The synthesis of cmnm^5^U nucleoside including reductive amination has been described before, however we found that the yields were poor when trying to reproduce the published synthesis pathway.^26^ Therefore, we developed a new synthetic pathway which proceeded reproducibly with much better yields. We decided to start with silylated 5-bromomethyl uridine **6** as a universal starting material for the synthesis of nm^5^U, mnm^5^U^7, 8^ and now cmnm^5^U (Scheme 2). We began the synthesis with 5-methyl uridine **4**, that is in the first place protected with (tBu)_2_Si and TBS protecting groups at the sugar hydroxy moiety (**5**).^7^ Subsequently, the 5-Me position was brominated using AIBN as an initiator and NBS, to enable the coupling of npe-protected glycine in the presence of base. These two steps were conducted without intermediate (**6**) purification steps. The formed secondary amine was protected with trifluoroacetyl group, to increase the stability of **7** during the column purification.^27^ The overall yield of these three steps was 43%. In the next step, the 3′and 5′position of the ribose were deprotected using HF in pyridine (**8**), followed by DMT-Cl protection of the primary alcohol providing compound **9**. Final product **10** was obtained after reaction with CED-Cl in the presence of base.

**Scheme 2.**
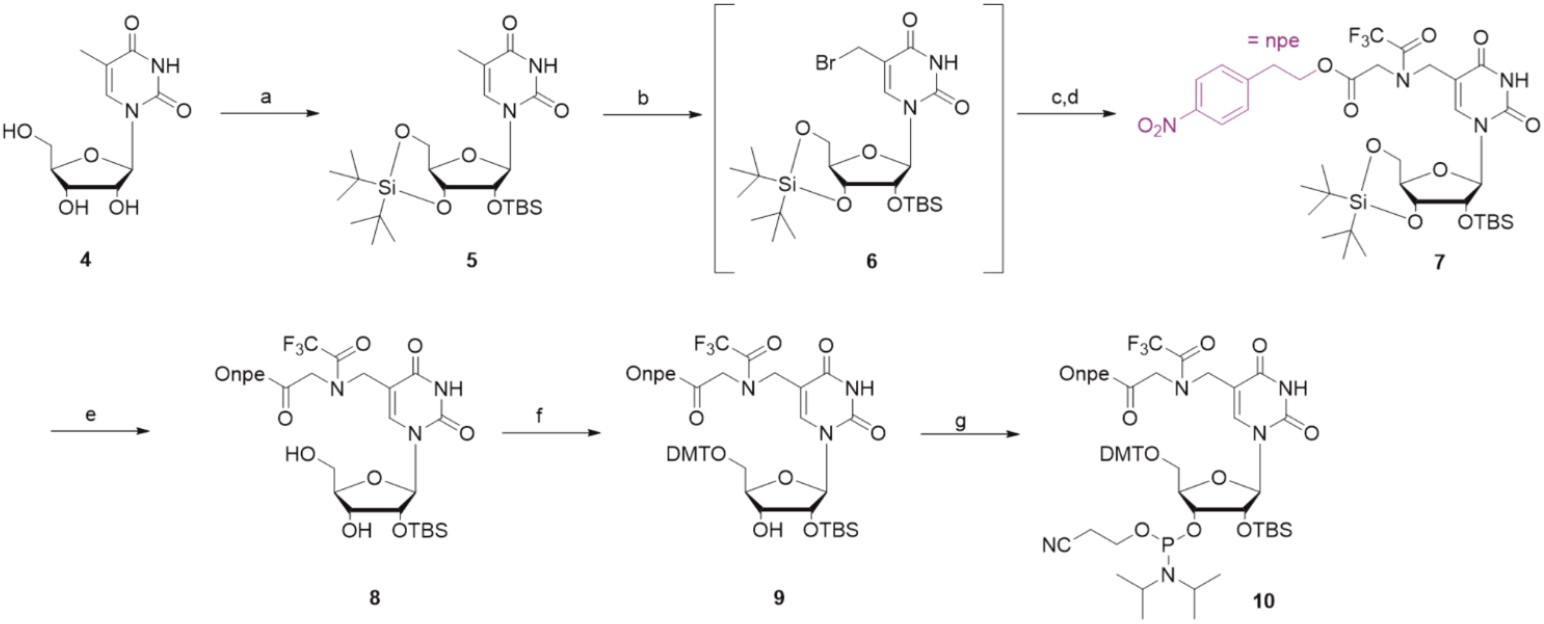
Reagents and conditions: a) (tBu)2Si(OTf)2, DMF, 0 °C - r.t., 1h; TBSCl, imidazole, 60°C, 1h, 88 %; b) NBS, AIBN, benzene 70 °C, 1.5h; c) HCl.glycine-Onpe, DIPEA, DMF, 0 °C - r.t.; d) TFAA, pyridine, 0 °C, 43 % over 3 steps; e) HF-pyridine, pyridine, DCM, 0°C, 1.5 h, 89 %; f) DMTCl, Py, 0°C to r.t. 73 %; g) CED-Cl, DIPEA, DCM 0°C to r.t. 70 %.

Both building blocks acp^3^U and cmnm^5^U phosphoramidites were incorporated into oligonucleotides using solid-phase oligonucleotide synthesis. First, we prepared two shorter strands **ON1** and **ON2** with incorporated modified phosphoramidite (X = acp^3^U; Y = cmnm^5^U) in the middle position, later we prepared longer oligonucleotide with acp^3^U incorporated at the 5’ end (**ON3**) and fully modified 2′-OMe oligonucleotide **ON4** with incorporated cmnm^5^U nucleoside at the 5’end (Fig. 3).

**Figure 3.**
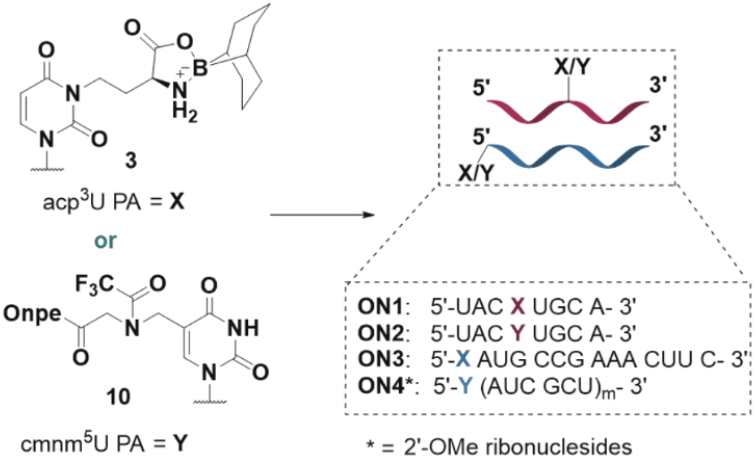
Oligonucleotide sequences with incorporated acp^3^U and cmnm^5^U.

The incorporation of both acp^3^U and cmnm^5^U proceeded without fluctuation of trityl values and comparably to the canonical nucleobases. Even during the synthesis of oligonucleotides **ON3** and **ON4** no decrease of the trityl value was observed and the phosphoramidites were stable during the prolonged time in solution. Neither double coupling nor an extension of the coupling time was needed, even though those conditions are often required for the insertion of modified nucleosides in order to obtain a decent yield. The most critical steps for a solid phase RNA synthesis are the deprotection steps, which could destroy the successfully incorporated non-canonical nucleosides. In case of cmnm^5^U strand, the npe protecting group was first deprotected on the beads using DBU/THF mixture (1:9) at room temperature for 2 hours. These deprotecting conditions did not cause any damage to the structure of the strands. Deprotection and cleavage of the oligonucleotides from the CPG-solid support material was achieved using aqueous AMA solution (NH_4_OH:methylamine 1:1) at 65 °C for 10 min, which simultaneously deprotected canonical nucleobases and homoserine moiety and cleaved the CF_3_CO group of cmnm^5^U strands. Following this second deprotection step was accomplished with TEA^x^3HF (65 °C, 1.5h). No other deprotection steps were required. All oligonucleotides were purified using reverse phase HPLC and the purity and structural integrity were evaluated by HPLC and MALDI-TOF as depicted in Fig. 4.

**Figure 4.**
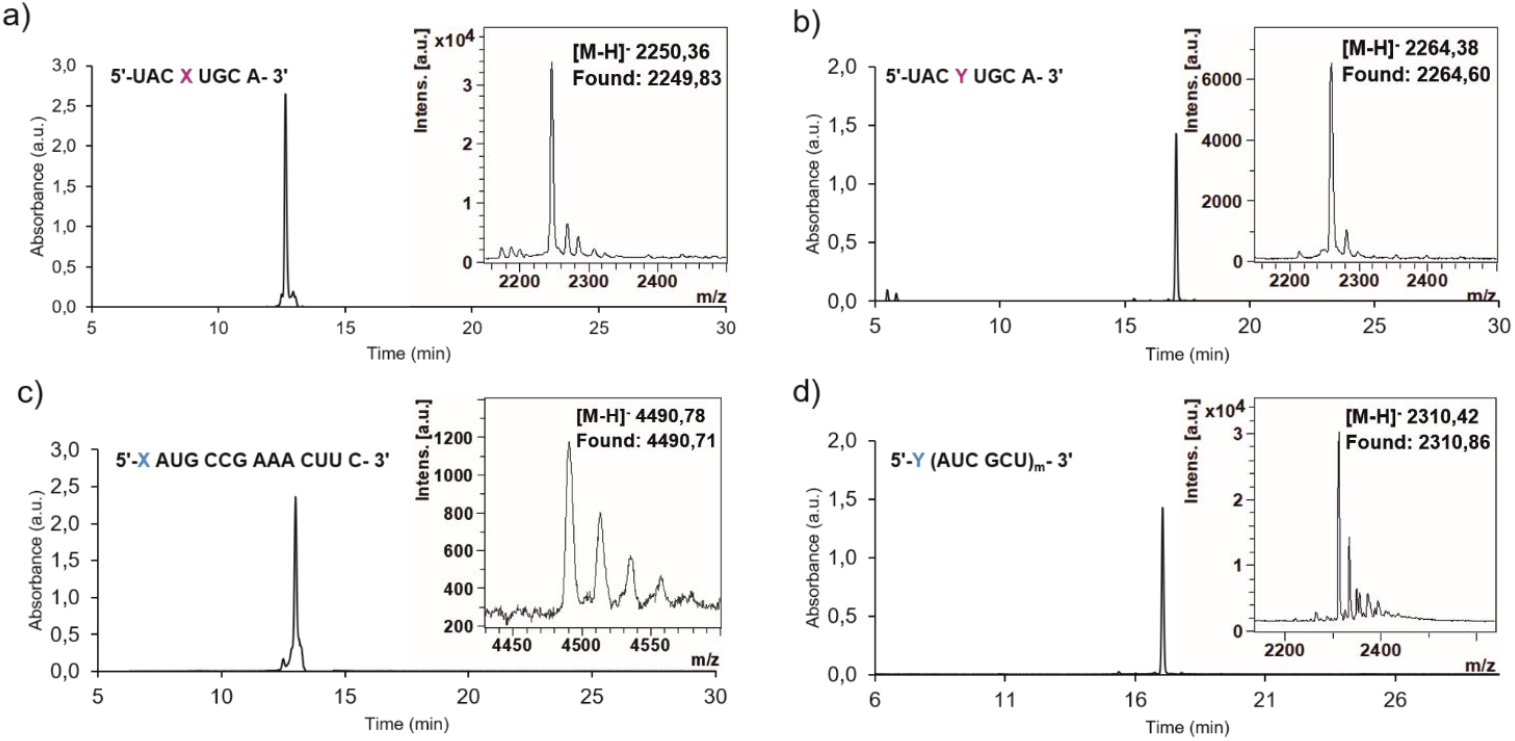
HPL chromatograms of purified oligonucleotides a) **ON1**, b) **ON2**, c) **ON3** and d) **ON4** with inserted MALDI-TOF mass spectrum.

## Conclusion

Hypermodified nucleosides have been recently extensively investigated for the unique properties connected to RNAs stability and function. Yet the emerging field of epitranscriptomics is still full of knowledge to be discovered. Especially acp^3^U and its involvement in native immunity and identification of enzymes responsible for its glycosylation are hot topic. For these purposes, oligonucleotides bearing acp^3^U are of a great interest. In our study, we showed elegant, simple and high yielding synthetic preparation of acp^3^U phosphoramidite in gram scale and its incorporation into various oligonucleotides. Moreover, we presented alternative synthesis of cmnm^5^U from universal starting material, that could be easily by split methodology converted not only to cmnm^5^U, but also nm^5^U and mnm^5^U simultaneously.

## Supporting Information

The data that support the findings of this study are available in the supplementary material of this article.^25^

## Acknowledgements

We thank the Deutsche Forschungsgemeinschat for supporting this research through the DFG grants: CA275/11-3 (ID:326039064), CRC1309 (ID: 325871075, A4), CRC1032 (ID:201269156, A5) and CRC1361 (ID: 393547839, P2). This project has received funding from the European Research Council (ERC) under the European Union’s Horizon 2020 Research and Innovation Program under grant agreement No. 741912 (EPiR). The research was supported by the BMBF in the framework of the Cluster4Future program (Cluster for Nucleic Acid Therapeutics Munich, CNATM) (Project ID: 03ZU1201AA).

## Entry for the Table of Contents

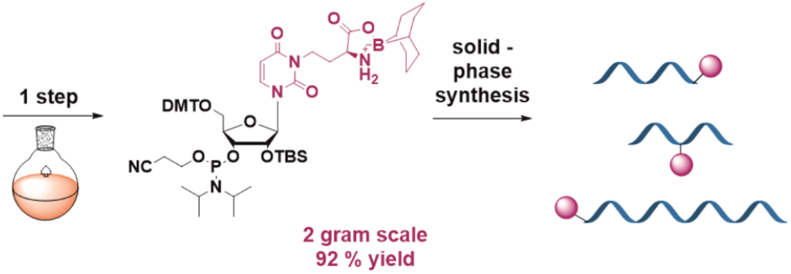

One-step, highly efficient and scalable synthesis of acp^3^U phosphoramidite is described as well as a new alternative synthesis of cmnm^5^U phosphoramidite from a universal intermediate. Latter could be scaled-up, split and aliquots turned simultaneously into cmnm^5^U, nm^5^U and mnm^5^U. The synthesis responds to the high demand of hypermodified nucleosides building blocks for the oligonucleotide synthesis.

