## Supplementary material for "Towards One-Step and Universal Synthesis of Hypermodified Uracil Nucleosides: acp^3^U and cmnm^5^U"

### Supporting Information

  **2.2 Synthesis of cmnm5U phosphoramidite** ……………………………………………………………….4

**3. Synthesis and characterization of oligonucleotides**…………………………………………………………..9

### 1. General information and instruments for nucleosides and phosphoramidites

Reagents were purchased from commercial suppliers and used without further purification unless otherwise stated. Anhydrous solvents, stored under inert atmosphere, were also purchased. All reactions involving air/moisture sensitive reagents/intermediates were performed under inert atmosphere using oven-dried glassware. Routine $^{1}$H NMR, $^{13}$C{$^{1}$H} NMR and $^{31}$P{$^{1}$H} NMR spectra were recorded on a Bruker Ascend 400 spectrometer (400 MHz for $^{1}$H NMR, 100 MHz for $^{13}$C NMR and 162 MHz for $^{31}$P NMR), Bruker Ascend 500 spectrometer (500 MHz for $^{1}$H NMR, 125 MHz for $^{13}$C NMR and 202 MHz for $^{31}$P NMR) or Bruker ARX 600 spectrometer (600 MHz for $^{1}$H NMR, 150 MHz for $^{13}$C NMR and 243 MHz for $^{31}$P NMR). Deuterated solvents used are indicated in the characterization and chemical shifts ($\delta$) are reported in ppm. Residual solvent peaks were used as reference.[1] All NMR $J$ values are given in Hz. COSY, HMQC and HMBC NMR experiments were recorded to help with the assignment of $^{1}$H and $^{13}$C signals. NMR spectra were analyzed using MestReNova software version 10.0. High Resolution Mass Spectra (HRMS) were measured on a Thermo Finnigan LTQ-FT with ESI as ionization mode. IR spectra were recorded on a Perkin-Elmer Spectrum BX II FT-IR instrument or Shimadzu IRSpirit FT-IR instrument. Both equipped with an ATR accessory. Column chromatography was performed with technical grade silica gel, 40-63 µm particle size. Reaction progress was monitored by Thin Layer Chromatography (TLC) analysis on silica gel 60 F254 and stained with *para*-anisaldehyde, potassium permanganate or cerium ammonium molybdate solution.

### 2. Synthesis and characterization data of phosphoramidites

#### 2.1 Synthesis of acp3U phosphoramidite

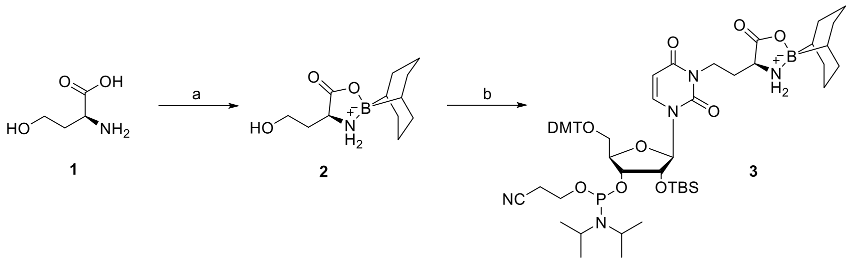

**Scheme S1:** Synthesis of acp$^{3}$U phosphoramidite ***3.*** Reagents and conditions: a) 9-BBN, MeOH, THF, 80 %; b) 2´-OTBS uracil phosphoramidite, PPh$_3$, DIAD, 1,4-dioxane, 0 °C-r.t., 3h, 92 %.

*(S)-4'-(2-hydroxyethyl)-9l4-boraspiro[bicyclo[3.3.1]nonane-9,2'-[1,3,2]oxazaborolidin]-5'-one* **2**

Title compound **2** was prepared according to previously published procedure. Analytical data agree with previously published procedure.[25] The reaction was conducted according to a published procedure with minor modifications. 1 L-homoserine (2) (0.371 g, 3.12 mmol) was suspended in methanol (25 mL) and heated under reflux until the mixture became clear. Then, a solution of 9-BBN, 9-borabicyclo(3.3.1)nonane (6.7 mL, 3.35 mmol) in tetrahydrofuran (0.5 M) was added dropwise. The reaction mixture was refluxed for 3 hours under inert atmosphere. The reaction mixture was concentrated and the crude product purified by silica gel chromatography eluting with 50% ethyl acetate-hexane to 100% ethyl acetate. The 9- BBN protected L-homoserine (3b) was obtained as a white solid (yield 80%); mp 112 – 115 o C. 1 H NMR (400 MHz, DMSO-d6) δ 0.49 (d, J = 0.5 Hz, 2H), 1.33 – 1.83 (m, 13H), 1.94 – 2.01 (m, 1H), 3.58 – 3.67 (m, 3H), 4.80 (t, J = 4.8 Hz, 1H), 5.86 – 5.91 (m, 1H), 6.41 – 6.46 (m, 1H); 13C NMR (101 MHz, DMSO-d6) δ 23.9, 24.4, 30.9, 31.3, 33.1, 52.3, 57.9, 174.0; HRMS (ESI): calculated for $C_{12}H_{23}BNO_3^+$ [M + H]+ : 240.1771; found 240.1764

*(2R,3R,4R,5R)-2-((bis(4-methoxyphenyl)(phenyl)methoxy)methyl)-4-((tert-butyldimethylsilyl)oxy)-5-(2,4-dioxo-3-(2-((R)-5'-oxo-9l4-boraspiro[bicyclo[3.3.1]nonane-9,2'-[1,3,2]oxazaborolidin]-4'-yl)ethyl)-3,4-dihydropyrimidin-1(2H)-yl)tetrahydrofuran-3-yl (2-cyanoethyl) diisopropylphosphoramidite* **3**

Uridine phosphoramidite (1.79g, 1 eq), BBN-protected homoserine **2** (0.624g, 1.2 eq) and PPh$_3$ (0.68g, 1.2 eq) were placed in round bottom flask and the flask was degassed and refilled with nitrogen. Dry 1,4-dioxane was added (15 mL) and the mixture was cooled to 0 °C before DIAD (0.52 mL, 1.2 eq) was added drop-wise. The mixture was allowed to warm up to r.t. and stirred for 3 hours. After completion of the reaction, the mixture was diluted with EtOAc, washed with NaHCO$_3$ and the organic phase was dried over sodium sulphate and evaporated. Residue was purified by column chromatography. Final product **3** was isolated in two fractions as separated isomers, both were freeze-dried from benzene.

**3:** Yield = 2.079 g (92%). Rf = 0.4 (1:1 iHex/EtOAc). $^{31}$P{$^{1}$H} NMR (202 MHz with cryoprobe, acetone-$d_6$, 298 K): $\delta$ (ppm) = 148.92; 150.16. HRMS (ESI) *m/z*: [M+H]$_+$ Calcd for $C_{57}H_{82}BN_5O_{11}PSi$ 1082,56053; Found 1082,56102.

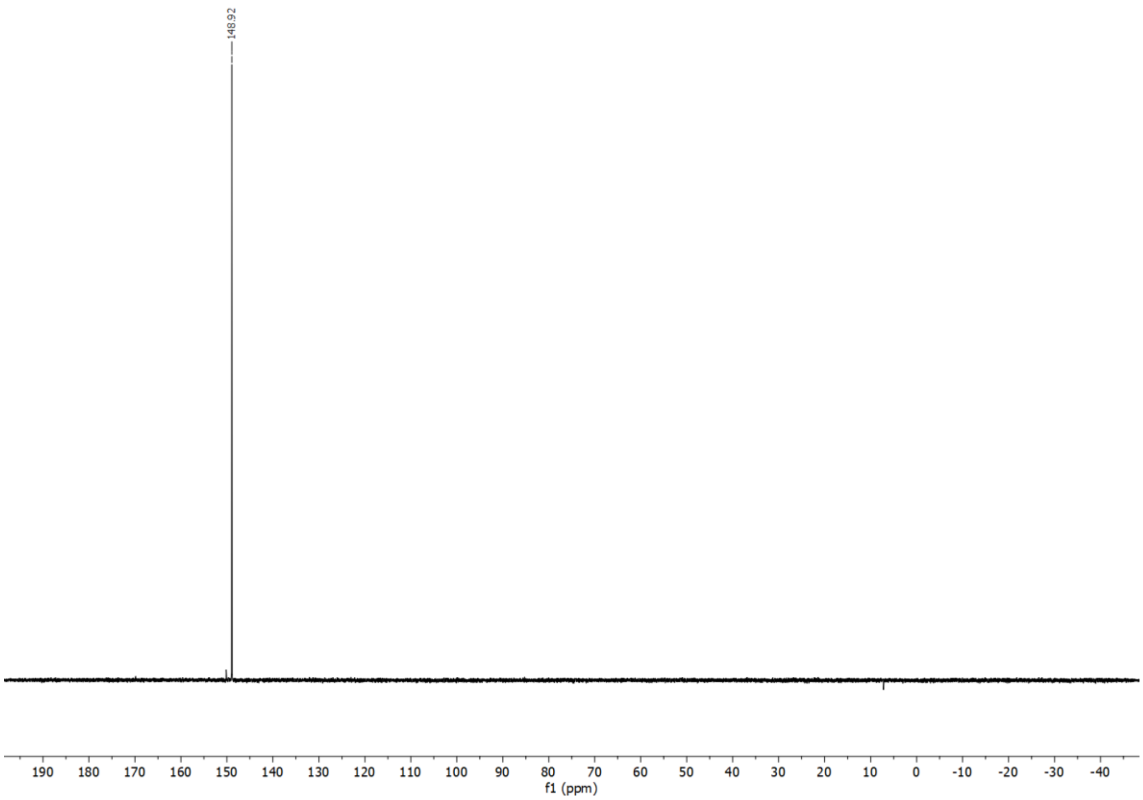
148.92
190 180 170 160 150 140 130 120 110 100 90 80 70 60 50 40 30 20 10 0 -10 -20 -30 -40
f1 (ppm)

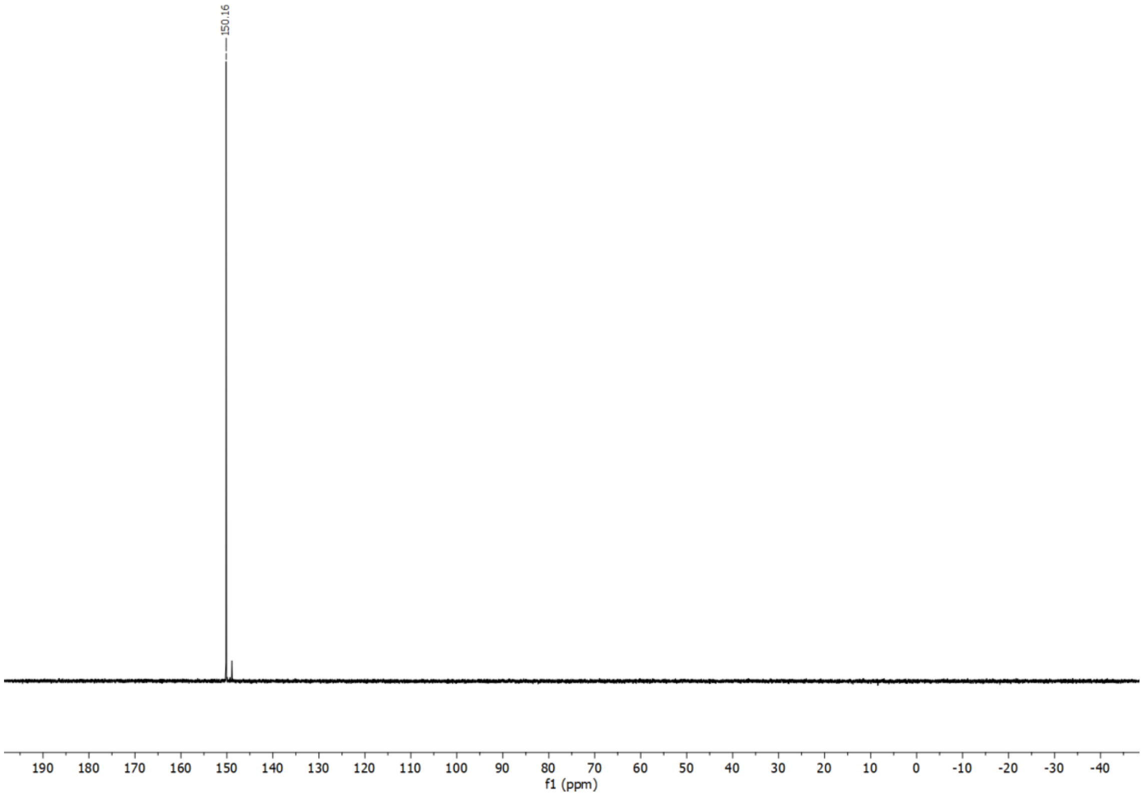
150.16
190 180 170 160 150 140 130 120 110 100 90 80 70 60 50 40 30 20 10 0 -10 -20 -30 -40
f1 (ppm)

### 2.2 Synthesis of cmnm5U phosphoramidite

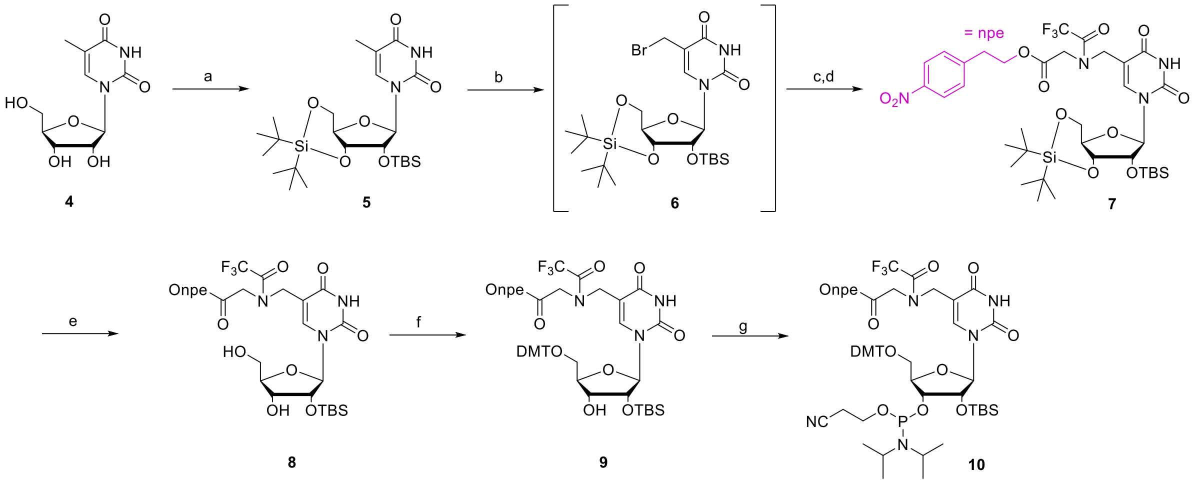

**Scheme S1**. Reagents and conditions: a) (tBu)2Si(OTf)2, DMF, 0 °C - r.t., 1h; TBSCl, imidazole, 60°C, 1h, 88 %; b) NBS, AIBN, benzene 70 °C, 1.5h; c) HCl.glycine-Onpe, DIPEA, DMF, 0 °C - r.t.; d) TFAA, pyridine, 0 °C, 43 % over 3 steps; e) HF-pyridine, pyridine, DCM, 0°C, 1.5 h, 89 %; f) DMTCl, Py, 0°C to r.t. 73 %; g) CED-Cl, DIPEA, DCM 0°C to r.t. 70 %.

**General procedure for the synthesis of compound 7:** Step 1. Compound **5** (0.5 g, 1 equiv.) was dissolved in dry benzene and degassed. To the solution NBS (0.24g, 1,3 eq) was added and the reaction was warmed to 60 °C. Then AIBN (0.08g, 0.5 eq) was added in one portion. The reaction was heated up to 70 °C and stirred for 2 hours. The TLC showed completion of the reaction. The reaction mixture was evaporated and used without purification. HCl.Glycine-Onpe (0.25g, 1 eq) was dissolved in dry DMF (4 mL) and TEA (0.28mL, 2eq) and the intermediate 6 was added dropwise as a solution in dry DMF. The reaction mixture was stirred overnight. Them, the mixture was diluted with EtOAc and washed with water and brine. The organic phase was dried over sodium sulfate and evaporated to dryness. In the last step, the crude was dissolved in dry pyridine (5mL) and cooled to 0 °C. TFAA (0.68mL, 5 eq) was added slowly and the mixture was stirred at 0 °C for 1h. The reaction was carefully poured into sat. $NaHCO_2$ and extracted with EtOAc. The organic phase was dried over sodium sulfate, evaporated and purified by column chromatography affording the product as a yellow foam.

**7:** Yield = 347 mg (43%). Rf = 0.3 (3:1 iHex/EtOAc). $^1$H NMR (500 MHz with cryoprobe, acetone-$d_6$, 298 K): *δ* (ppm) = 10.33 (s, 0.7H); 10.06 (s, 0.3H); 8.21 – 8.17 (m, 2H); 7.76 (s, 0.4H); 7.68 (s, 0.2H); 7.62 – 7.58 (m, 2.4H); 5.76 (d, *J* = 17.7 Hz, 0.5H); 5.70 (s, 0.5H); 4.55 – 4.40 (m, 6H); 4.26 – 4.07 (m, 5H); 3.19 – 3.12 (m, 2H); 1.07 – 1.04 (m, 18H); 0.96 – 0.94 (m, 9H); 0.22 – 0.18 (m, 3H); 0.18 – 0.15 (m, 3H). $^{13}$C NMR (126 MHz, Acetone) δ 168.28, 163.62, 149.62, 146.17, 130.20, 123.43, 108.06, 93.33, 75.74, 75.28, 74.81, 67.30, 65.08, 64.76, 50.02, 49.99, 48.76, 48.72, 34.36, 29.41, 29.25, 29.10, 28.94, 28.79, 28.64, 28.48, 27.00, 26.61, 25.45, -4.95, -5.61. HRMS (ESI) *m*/*z*: $[M+H]^+$ Calcd for C36H54F3N4O11Si2 831,32742; Found 831,3265.

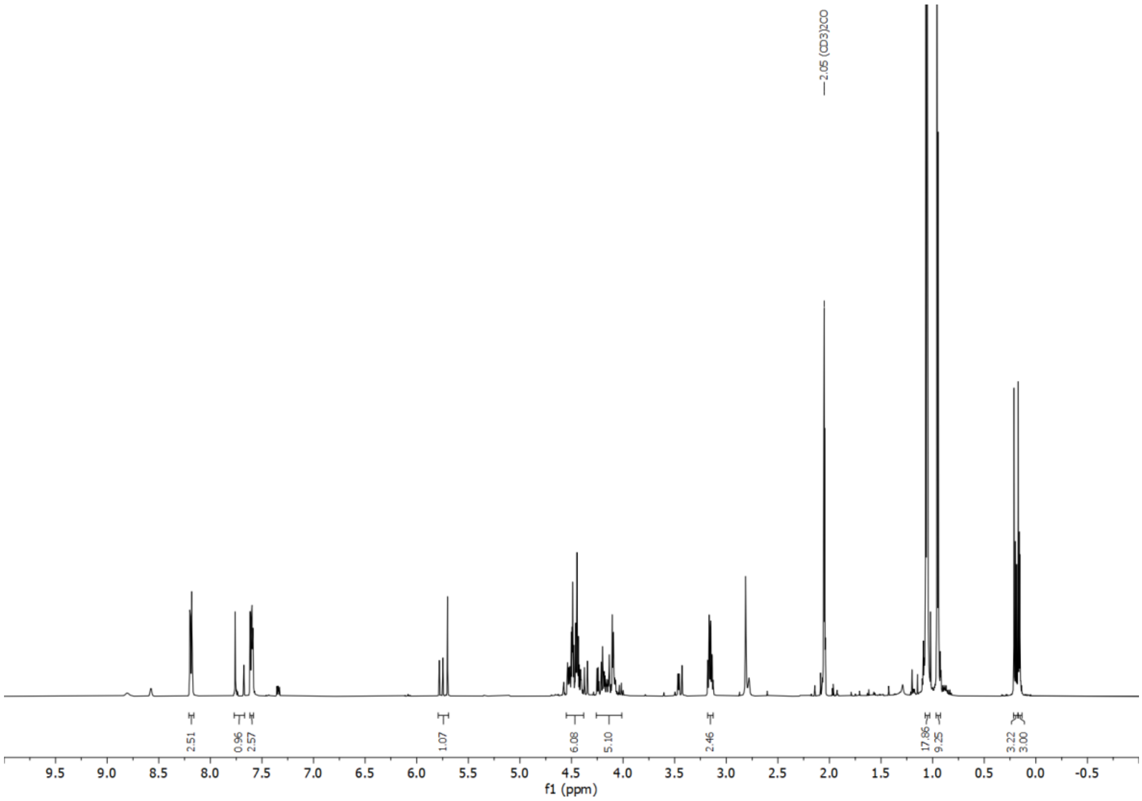

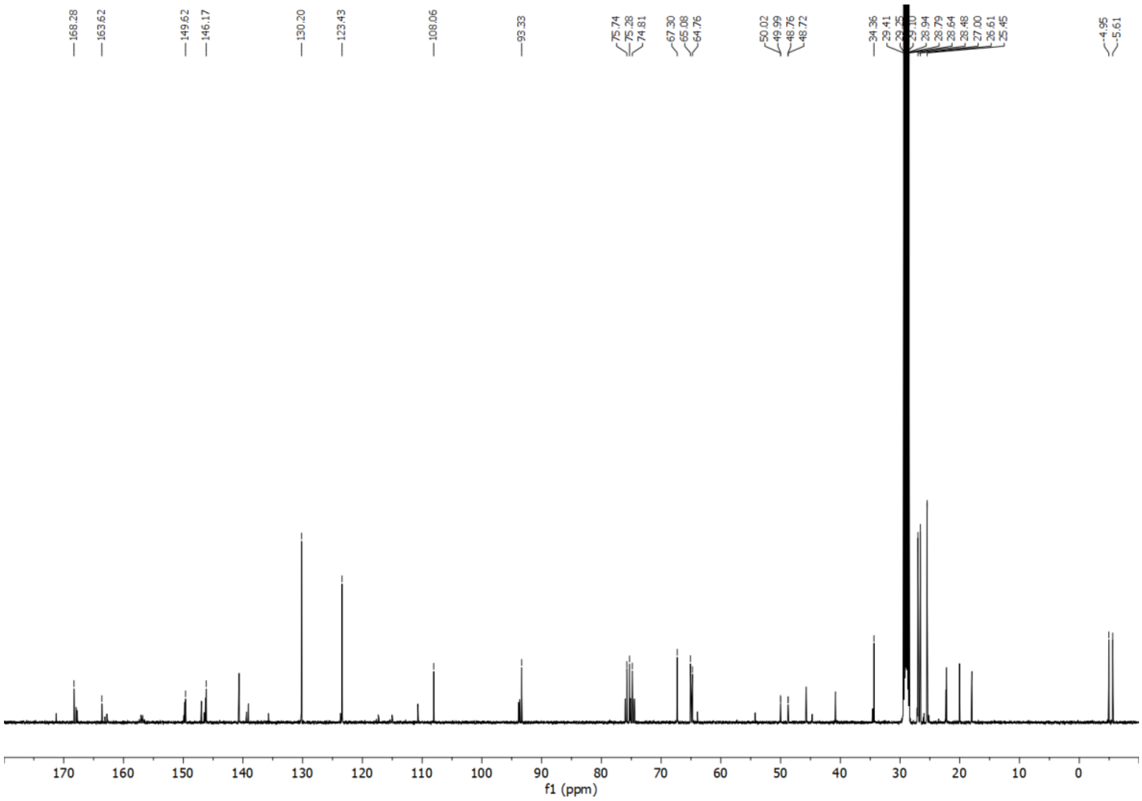

**General procedure for the synthesis of compound 8**

Compound **7** (1 equiv.) was added to a plastic flask and dissolved in dry 9:1 $CH_2Cl_2$/pyridine. The solution was stirred at 0°C. Finally, HF•pyridine (0.06uL, from a commercial solution containing 70% HF and 30% pyridine) was added and the reaction was stirred at 0°C for 2 h. After that, the reaction was quenched with aqueous saturated $NaHCO_3$ and $CH_2Cl_2$ was added. The organic layer was separated and the crude was further extracted with $CH_2Cl_2$. The combined organic layers were dried ($Na_2SO_4$), filtered and concentrated. The crude was purified by silica gel column chromatography affording the product as a white foam. The product is a mixture of rotamers (as seen in the NMR spectrum).

**8**: Yield 270 mg (89 %). Rf = 0.6 (2:3 iHex/EtOAc). $^1$H NMR (500 MHz with cryoprobe, DMSO-$d_6$, 298 K): $\delta$ (ppm) = 11.57 (d, $J$ = 9.9 Hz, 1H); 8.16 – 8.12 (m, 2H); 8.04 (s, 0.5H); 7.97 (s, 0.5H); 7.55 – 7.51 (m, 2H); 5.78 (dd, $J$ = 11.7 Hz, $J$ = 5.0 Hz, 1H); 5.17 (dt, $J$ = 29.7 Hz, $J$ = 5.0 Hz, 1H); 5.03 (dd, $J$ = 8.1 Hz, $J$ = 5.4 Hz, 1H); 4.45 – 4.20 (m, 5H); 4.18 – 4.08 (m, 3H); 3.98 – 3.87 (m, 2H); 3.04 (dt, $J$ = 9.1 Hz, $J$ = 6.4 Hz, 2H); 0.80 (d, $J$ = 7.5 Hz, 9H); 0.01 – -0.05 (m, 6H). $^{13}$C{$^1$H} NMR (125 MHz with cryoprobe, DMSO-$d_6$, 298 K): $\delta$ (ppm) = 168.4; 167.8; 163.5; 162.9; 150.5; 150.4; 149.7; 146.5; 146.5; 146.4; 140.6; 138.6; 136.4; 130.4; 130.3; 124.2; 123.6; 108.2; 108.0; 88.4; 87.8; 85.5; 85.3; 76.1; 75.8; 70.1; 69.8; 65.1; 64.8; 61.0; 60.6; 60.0; 49.8; 48.7; 45.9; 45.1; 34.1; 25.8; 25.7; 18.0; 18.0; -4.7; -4.8; -5.1; -5.2. HRMS (ESI) $m/z$: $[M+H]^+$ Calcd for $C_{28}H_{38}F_3N_4O_{11}Si$ 691,2253; Found 691,2246.

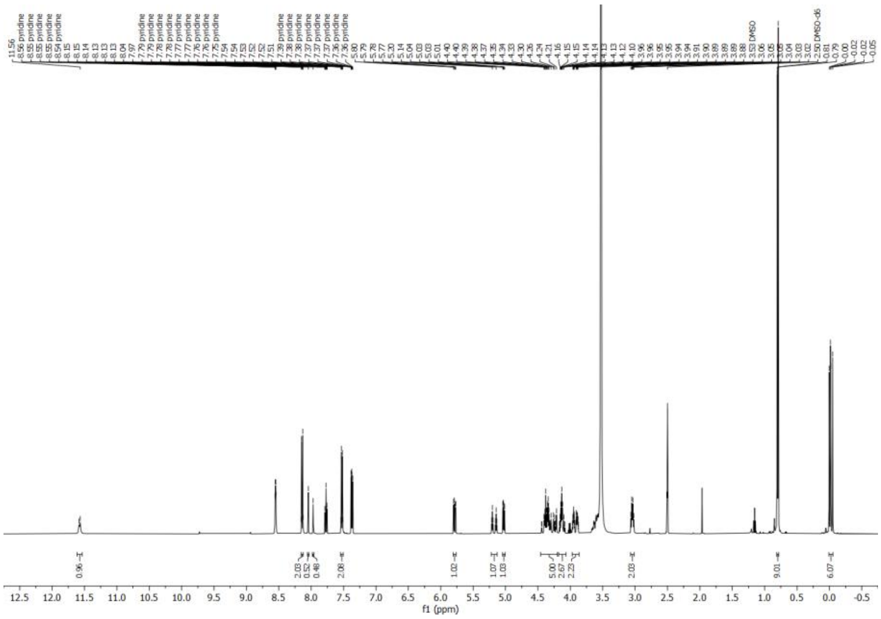

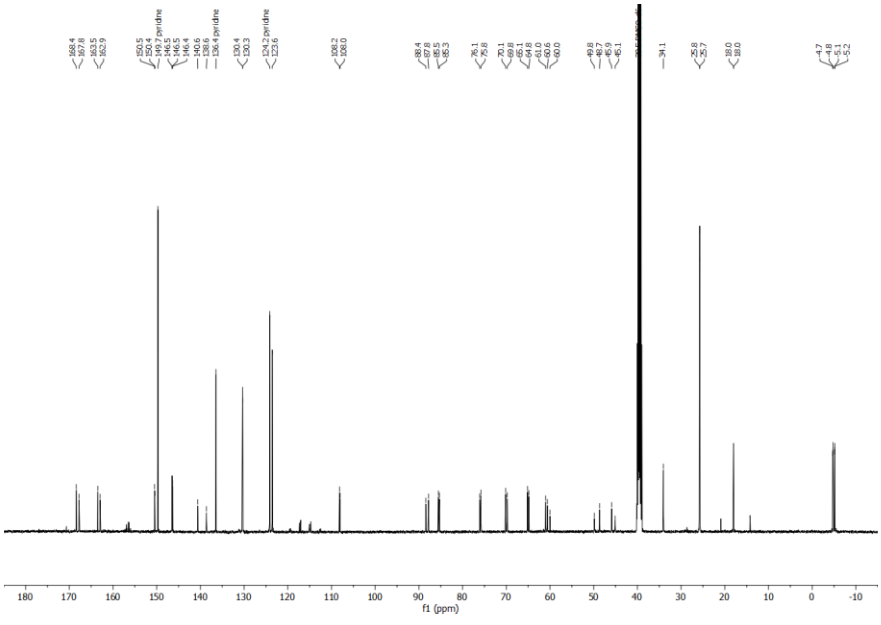

**General procedure for the synthesis of compound 9:** Compound **8** (1 equiv.) was dissolved in dry pyridine (4 mL) and stirred under nitrogen atmosphere at r.t. 4,4-Dimethoxytrityl chloride (0.2g, 1.5 equiv.) was added in one portion and the reaction was stirred at r.t. overnight. After that, the crude was quenched with MeOH and diluted with EtOAc. EtOAc solution was washed with water, dried over sodium sulphate, concentrated and purified by silica gel column chromatography (eluent containing 0.1% pyridine) affording the product as a white foam.

**9**: Yield 289 mg (73 %). Rf = 0.3 (2:1 iHex/EtOAc). $^{1}$H NMR (500 MHz with cryoprobe, acetone-$d_6$, 298 K): $\delta$ (ppm) = 10.35 (s, 1H); 8.17 (dq, $J$ = 8.7 Hz, $J$ = 2.3 Hz, 2H); 7.91 (s, 0.7H); 7.71 (s, 0.3H); 7.55 (ddd, $J$ = 8.9 Hz, $J$ = 4.2 Hz, $J$ = 2.9 Hz, 4H); 7.46 – 7.42 (m, 3H); 7.38 – 7.30 (m, 4H); 7.26 – 7.21 (m, 1H); 6.89 (td, $J$ = 8.5 Hz, $J$ = 7.9 Hz, $J$ = 1.3 Hz, 4H); 5.89 (d, $J$ = 4.0 Hz, 0.7H); 5.84 (d, $J$ = 3.3 Hz, 0.3H); 4.53 – 4.31 (m, 5H); 4.23 – 4.09 (m, 3H); 4.03 – 3.84 (m, 3H); 3.78 (d, $J$ = 1.9 Hz, 6H); 3.50 (dd, $J$ = 10.9 Hz, $J$ = 4.8 Hz, 0.8H); 3.41 (dd, $J$ = 10.9 Hz, $J$ = 2.5 Hz, 1.2H); 3.14 (t, $J$ = 6.5 Hz, 1.5H); 3.07 (t, $J$ = 6.6 Hz, 0.5H); 0.92 (d, $J$ = 2.0 Hz, 9H); 0.13 (d, $J$ = 2.6 Hz, 6H). $^{13}$C{$^{1}$H} NMR (125 MHz with cryoprobe, acetone-$d_6$, 298 K): $\delta$ (ppm) = 169.2; 164.4 (163.4); 159.6 (159.7); 150.9; 150.7; 147.8; 147.1; 146.1; 141.9; 136.8; 136.8; 136.6; 131.1 (131.0); 129.1 (129.0); 128.7 (128.7); 127.6 (127.7); 124.6; 124.3 (124.3); 114.0; 114.0; 114.0; 113.9; 109.2; 91.5; 90.3; 87.3 (87.2); 84.3 (83.9); 76.8 (76.5); 71.3; 65.9 (65.6); 64.5; 55.5; 51.1; 48.8; 47.4; 35.2; 35.2; 26.2 (25.2); 18.8; 18.7; -4.6; -4.7. HRMS (ESI) $m/z$: $[M+H]^+$ Calcd for $C_{49}H_{56}F_3N_4O_{13}Si$ 993,35598; Found 993,35489.

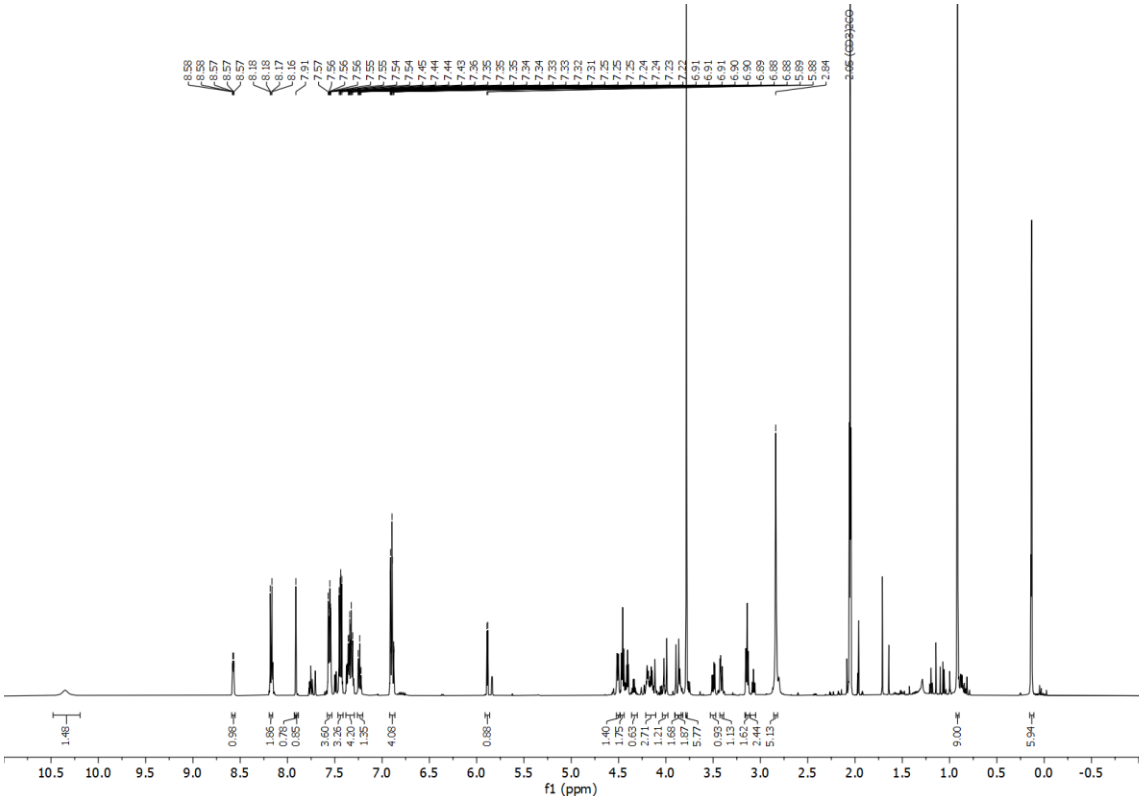

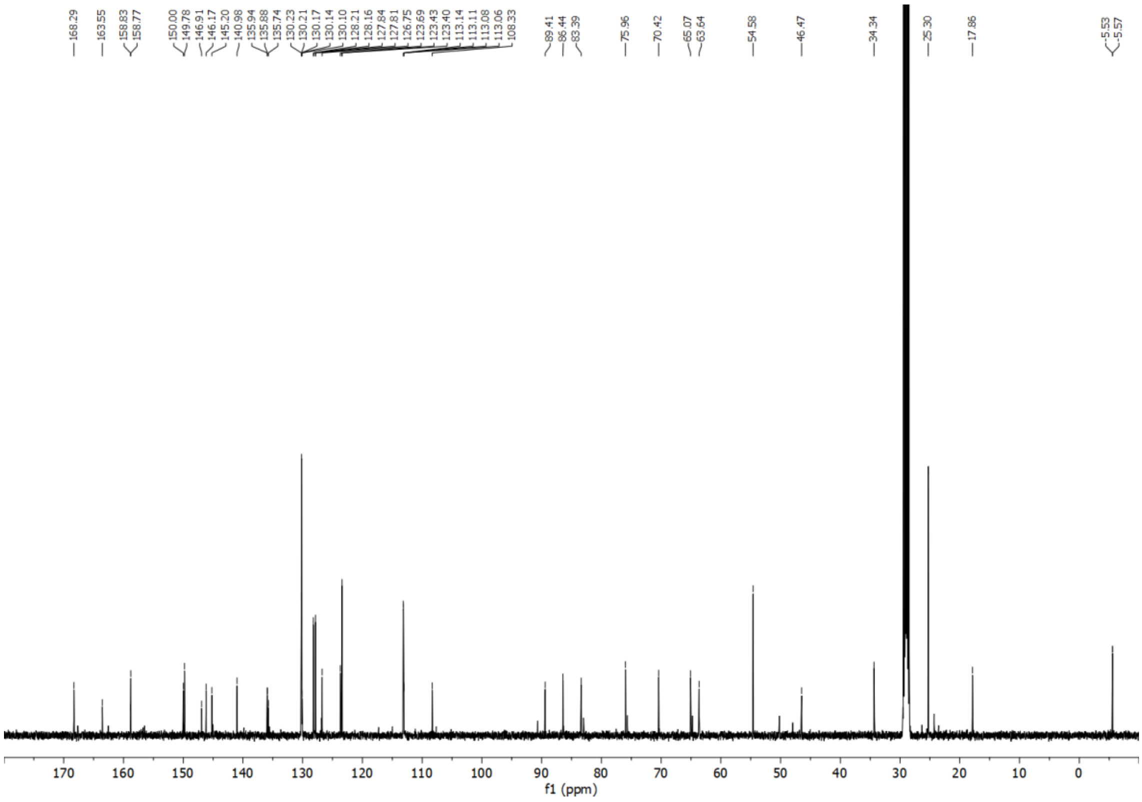

**General procedure for the synthesis of compound 10:** Compound **9** (0.17g, 1 equiv.) was added to a dry-oven round-bottom flask and dissolved in dry $CH_2Cl_2$ (3 mL). The solution was stirred under Argon atmosphere at 0°C. DIPEA (0.13uL, 4 equiv.) was added dropwise. Finally, 2-cyanoethyl *N*,*N*-diisopropylchlorophosphoramidite (0.1uL, 2.5 equiv.) was added dropwise. The reaction was stirred at r.t. for 3 h. After that, the reaction was stopped and diluted with $CH_2Cl_2$. The crude was washed with aqueous saturated $NaHCO_3$ and the organic layer was separated. The crude was further extracted with $CH_2Cl_2$. The combined organic layers were dried ($Na_2SO_4$), filtered and concentrated. The crude was purified by silica gel column chromatography (eluent containing 0.1% pyridine).

The products were isolated as a mixture of diastereoisomers as a white foam. Finally, the product was lyophilized from benzene.

**10:** Yield = 143 mg (70%). Rf = 0.6 (2:1 iHex/EtOAc). $^{31}P\{^{1}H\}$ NMR (202 MHz with cryoprobe, acetone-$d_6$, 298 K): *δ* (ppm) = 149.9 (149.3); 149.2 (149.2). HRMS (ESI) *m/z*: $[M+H]^+$ Calcd for $C_{58}H_{73}F_3N_6O_{14}PSi$ 1193,46383; 1193,46471.

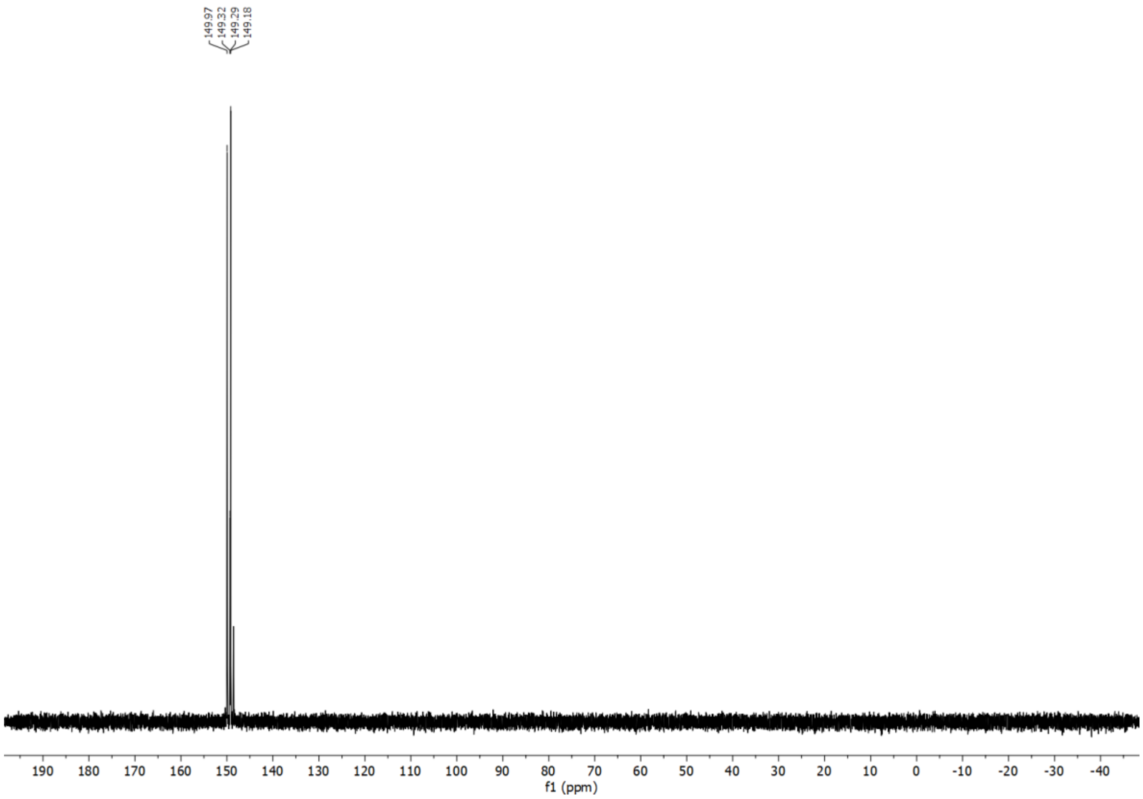

**3. General information and instruments for oligonucleotides**

**3.1. Synthesis and purification of oligonucleotides**

Phosphoramidites of 2'-O-Me ribonucleosides (2'-OMe-Bz-A-CE, 2'-OMe-Dmf-G-CE, 2'-OMe-Ac-C-CE and 2'-OMe-U-CE) and deoxyribonucleosides (Bz-dA-CE, Dmf-dG-CE, Ac-dC-CE, T-CE) were purchased from LinkTech and Sigma-Aldrich. Oligonucleotides (ONs) were synthesized on a 1 µmol scale using RNA SynBaseTM CPG 1000/110 and High Load Glen UnySupportTM as solid supports using an RNA automated synthesizer (Applied Biosystems 394 DNA/RNA Synthesizer) with a standard phosphoramidite chemistry. ONs were synthesized in DMT-OFF mode using DCA as a deblocking agent in CH2Cl2, BTT or Activator 42® as activator in MeCN, Ac2O as capping reagent in pyridine/THF and I2 as oxidizer in pyridine/H2O.

**3.2. Cleavage from beads and precipitation of the synthesized ON**

The solid support beads were suspended in a 1:1 aqueous solution mixture (0.6 mL) of 30% NH4OH and 40% MeNH2. The suspension was heated at 65°C (8 min for SynBaseTM CPG 1000/110 and 60 min for High Load Glen UnySupportTM). The ONs containing dipeptide-modified carbamoyl adenosine derivatives were cleaved from the solid support beads using a 30% $NH_4OH$ aqueous solution (0.6 mL) at r.t. overnight. Subsequently, the supernatant was collected, and the beads were washed with water (2×0.3 mL). The combined aqueous solutions were concentrated under reduced pressure using a SpeedVac concentrator. After that, the crude was dissolved in DMSO (100 µL) and the ON was precipitated by adding 3 M NaOAc in water (25 µL) and n-butanol (1 mL). The mixture was kept at -80°C for 2 h and centrifuged at 4°C for 1 h. The supernatant was removed, and the white precipitate was lyophilized.

**3.2. Purification of the synthesized ON by HPLC and desalting**

The crude was purified by semi-preparative HPLC (1260 Infinity II Manual Preparative LC System from Agilent equipped with a G7114A detector) using a reverse-phase (RP) VP 250/10 Nucleodur 100-5 C18ec column from

Macherey-Nagel (buffer A: 0.1 M AcOH/Et3N pH 7 in H2O and buffer B: 0.1 M AcOH/Et3N pH 7 in 20:80 $H_2O$/MeCN; Gradient: 2-40% of B in 45 min; Flow rate = 5 mL·min-1). The purified ON was analyzed by RP-HPLC (1260 Infinity II LC System from Agilent equipped with a G7165A detector) using an EC 250/4 Nucleodur 100-3 C18ec from Macherey-Nagel (Gradient: 2-40% of B in 30 min; Flow rate = 1 mL·min-1). Finally, the purified ON was desalted using a C18 RP-cartridge from Waters.

| Sequence | *tR (min)* | *m/z calcd. for [M-H]-* | *found* |
|---|---|---|---|
| **ON1**: 5'-UAC X UGC A- 3' | 12.649 | 2250,4 | 2249,8 |
| **ON2:** 5'-UAC Y UGC A- 3' | 12.474 | 2264,4 | 2264,6 |
| **ON3**: 5'-X AUG CCG AAA CUU C- 3' | 13.001 | 4490,8 | 4490,7 |
| **ON4*:** 5'-Y (AUC GCU)$_m$- 3' | 17.055 | 2310,4 | 2310,8 |

**ON1**

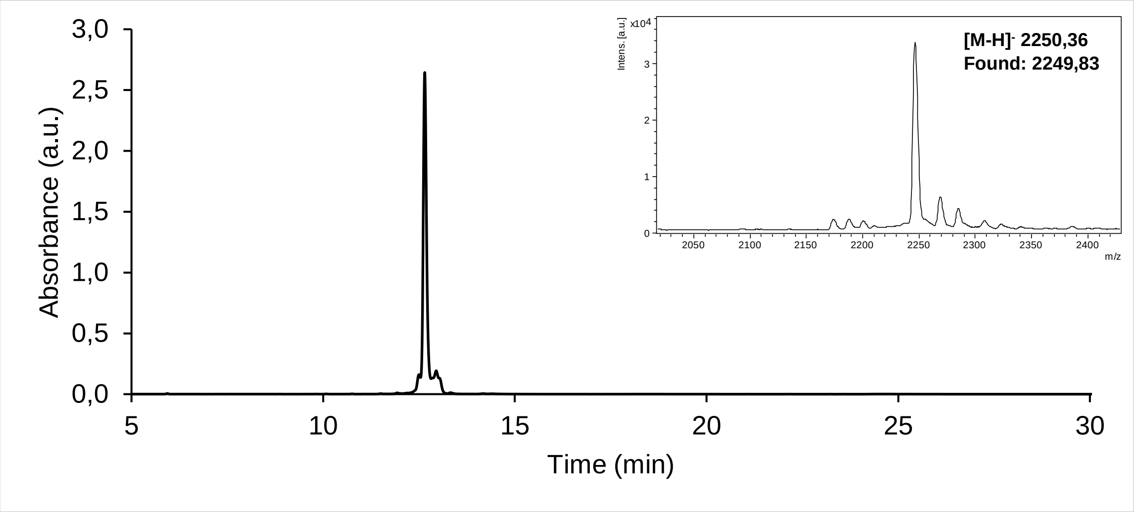

**ON2**

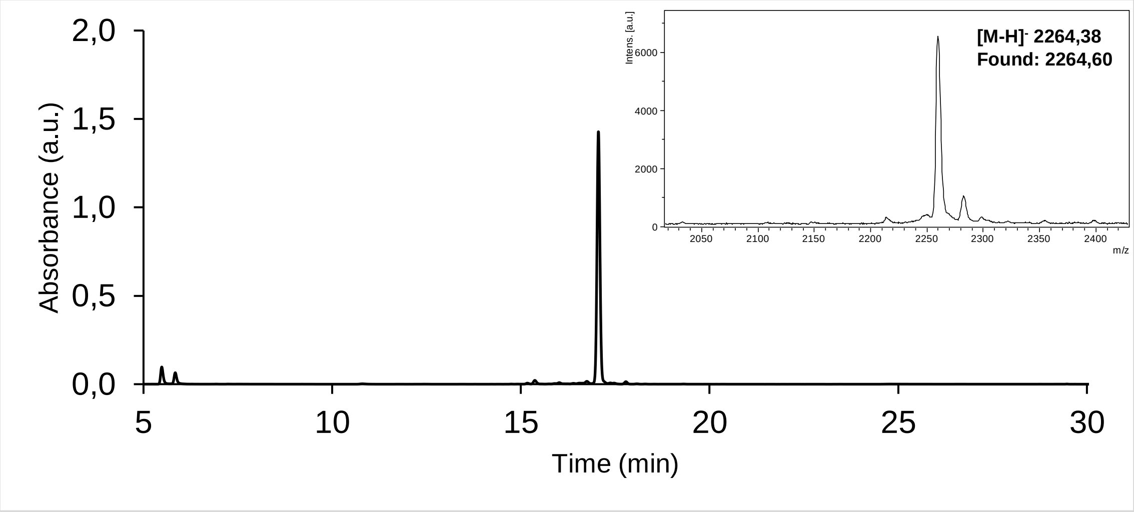

**ON3**

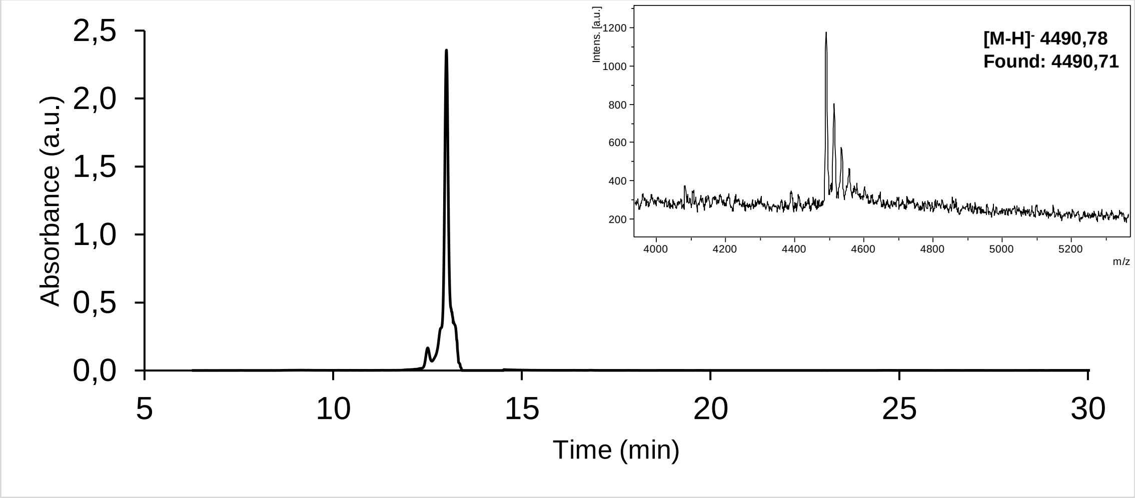
[M-H]- 4490,78
Found: 4490,71
Absorbance (a.u.)
Time (min)
Intens. [a.u.]
m/z

**ON4**

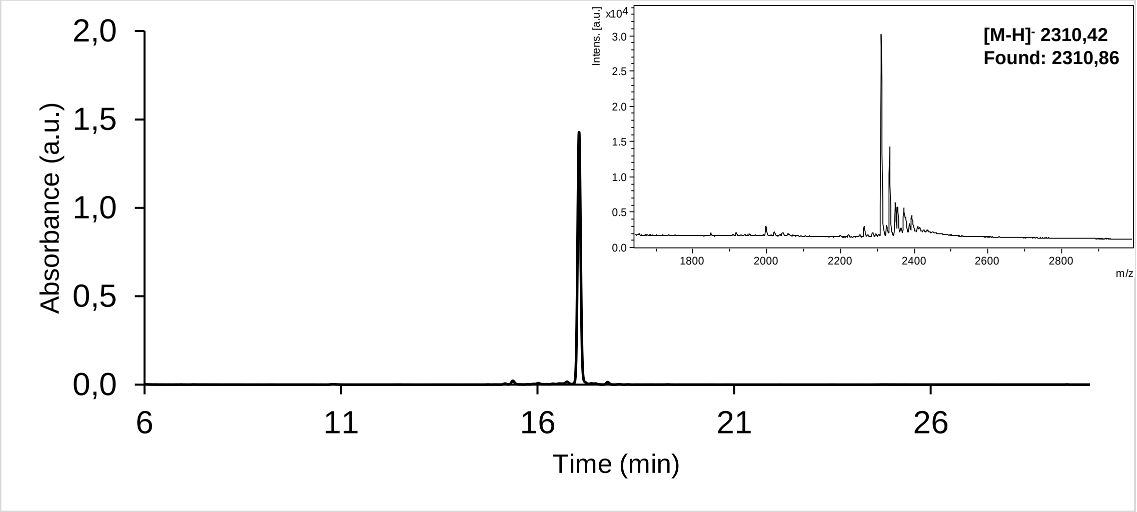
[M-H]- 2310,42
Found: 2310,86
Absorbance (a.u.)
Time (min)
Intens. [a.u.]
m/z
